# Reduced Plasma Ascorbic Acid Levels in Recipients of Myeloablative Conditioning & Hematopoietic Cell Transplantation

**DOI:** 10.1101/539007

**Authors:** Mahmood Rasheed, Gary Simmons, Bernard Fisher, Kevin Leslie, Jason Reed, Catherine Roberts, Ramesh Natarajan, Alpha Fowler, Amir Toor

**Affiliations:** Department of Medicine, Virginia Commonwealth University, Richmond, VA 23298.; Department of Physics, Virginia Commonwealth University, Richmond, VA 23298.

**Keywords:** Ascorbic acid, Parenteral Vitamin C, Hematopoietic cell transplantation, Endothelial injury, Graft versus host disease

## Abstract

Hematopoietic cell transplantation (HCT) conditioned using myeloablative conditioning (MAC) is complicated by end organ injury due to endothelial dysfunction and graft versus host disease. Mucositis and oxidant injury results in micronutrient deficiency. Ascorbic acid (AA) levels were measured in 15 patients undergoing HCT conditioned with MAC (11 allogeneic and 4 autologous HCT). Ascorbate levels declined post conditioning to 27.3 (±14.1) by day 0 (p <0.05 compared with baseline), reaching a nadir level of 21.5 (±13.8) on day 14 (p <0.05) post-transplant. Patients undergoing allogeneic HCT continued to have low AA levels to day 60 post transplant, whereas recipients of autologous HCT recovered plasma AA levels to normal. The role of AA in maintaining endothelial function and hematopoietic as well as T cell recovery is provided, developing the rationale for repletion of vitamin C following HCT.

## Introduction

Patients with hematologic malignancy undergoing stem cell transplant face competing hazards of non-relapse mortality (NRM) as well as relapsed malignancy following hematopoietic cell transplantation (HCT). Reduction in the intensity of conditioning therapy minimizes NRM, at the expense relapsed malignancy. Conditioning intensity, while important for relapse mitigation in some diseases (i.e., acute myelogenous leukemia) is associated with tissue injury, particularly endothelial injury that leads to endogenous tissue antigen presentation to donor T cells, thus, provoking graft versus host disease (GVHD) and contributing to NRM.^1,2,3,4^ Intense immunosuppression for mitigation of GVHD may diminish the *graft versus malignancy* response that may be expected from an immune-competent graft. Therefore, it is critical that therapy to mitigate endothelial and tissue injury in HCT recipients be developed to aid in balancing the competing hazards of tissue injury and malignancy relapse. One agent that may serve this purpose is ascorbic acid (vitamin C). Ascorbic acid is an effective anti-inflammatory agent with its demonstrated ability to inhibit NF-kB-driven inflammatory cytokines IL-6, IL-8, and TNF-α expression, and by its ability to enhance vasodilator nitric oxide (NO) production.^5,6^ Further, hypovitaminosis C is well described in critically ill patients. In this population, intravenous ascorbic acid increases nitric oxide bioavailability in the microcirculation. Vitamin C action through this mechanism limits endothelial dysfunction, lowers inflammatory biomarkers, improves organ dysfunction and thus improves clinical outcomes.^7^

Patients undergoing myeloablative allogeneic HCT often develop nutritional deficiencies in immediate post-transplant periods. Multiple factors, including conditioning regimen-related oxidant injury, gastrointestinal mucositis, and increased nutritional requirements due to catabolic stresses converge to place patients at risk for general malnutrition and micronutrient deficiencies. Malnutrition impacts clinical outcomes in patients undergoing HCT particularly in the presence of ongoing endothelial injury such as by administration of calcineurin inhibitors. This observation is particularly germane to the HCT population, in whom myeloablative conditioning results in a pro-inflammatory state characterized by elevated inflammatory cytokines. The resulting inflammatory milieu leads to endothelial injury and contributes to clinical syndromes such as sinusoidal obstruction syndrome, pulmonary alveolar hemorrhage, and increased infection risk. Over time the resulting inflammatory milieu increases antigen-presenting cell activity and likely contributes to subsequent development of GVHD. ^1^ Collectively these processes as described contribute to non-relapse mortality following allogeneic HCT. They also underpin the evolving interest in both the anti-inflammatory and endothelial protective properties of ascorbic acid.

It was hypothesized that patients undergoing myeloablative allogeneic SCT become deficient in micronutrients due to poor oral intake and mucosal injury, particularly ascorbic acid which is water soluble and rapidly metabolized in the prooxidative, pro-inflammatory state which follows conditioning therapy, particularly conditioning with irradiation and alkylator based regimens. Ascorbate administered in parenteral nutrition may not be adequate in this circumstance.^8^ Further, depletion of ascorbic acid may be caused by endothelial injury associated with GVHD prophylaxis regimens. To examine this possibility, the kinetics of plasma ascorbic acid were investigated in patients starting prior to the initiation of conditioning, and at multiple timepoints up to day 60 following myeloablative conditioning and HCT.

## Methods

Patients were enrolled prospectively in a Virginia Commonwealth University (VCU) Institutional Review Board (IRB)-approved study. Patients provided written informed consent prior to enrollment. Patients undergoing myeloablative conditioning followed by either HLA matched related or unrelated donor stem cell transplantation or those undergoing autologous stem cell transplant were eligible for participation. Blood samples to measure ascorbic acid levels were drawn prior to transplant and on days 0, 14, 30, and 60 post-SCT. Plasma ascorbic acid levels were determined via a modified fluorescence end-point assay following deproteinization as previously described.^9^ Patients were followed for development of mucositis and GVHD, with event severity coded by WHO grading criteria and Glucksberg grade, respectively.

## Results

### Plasma Ascorbic acid Levels

A total of 15 patients were enrolled from October 2015 through May 2016 (Table 1). Mean plasma ascorbic acid levels were 40.8 µmol/L (±18.4) (normal range: 50-80 µmol/L) at baseline before conditioning began. Plasma AA levels had fallen to 27.3 (±14.1) by day 0 (p <0.05 compared with baseline*) and reached a nadir at 21.5 (±13.8) on day 14 (p <0.05*) post-transplant (Figure 1). Ascorbic acid levels recovered to 34.2 (±20.5) (p=NS compared with baseline) by day 30, and 37.2 (±27.9) (p=NS compared with baseline) at day 60 following SCT (Figure 1).

**Table 1.** Patient characteristics and details of conditioning regimen (N=15).

|  |  |
| --- | --- |
| Male/Female | 6/9 |
| Age (range) | 50 years (31-65) |
| Transplant |  |
| *HLA matched related donor\** | 3 |
| *HLA matched unrelated donor\** | 8 |
| *Autologous* | 4 |
| Conditioning regimen** |  |
| *TBI+CY* | 1 |
| *Bu+CY* | 5 |
| *Bu+FLU* | 3 |
| *FLU+MEL* | 2 |
| *BEAM* | 3 |
| *MEL* | 1 |

Primary diagnoses: Myelofibrosis (2), acute myelogenous leukemia (2), acute lymphoblastic leukemia (2), myelodysplastic syndrome (3), non-Hodgkin lymphoma (3), Hodgkin lymphoma (1), multiple myeloma (1), and chronic myelogenous leukemia (1).
\*- HLA matching was at the allelic level
\*\*- Bu+CY- busulfan 0.8 mg/kg IV every 6 hours for 16 doses+cyclophosphamide 60 mg/kg daily for 2 doses; Bu+FLU- busulfan 130 mg/m<sup>2</sup> daily for 4 doses+fludarabine 30 mg/m<sup>2</sup> for four doses; FLU+MEL- single dose of melphalan 140 mg/m<sup>2</sup>+fludarabine 30 mg/m<sup>2</sup> for 4 doses; BEAM- single dose of carmustine 300 mg/m<sup>2</sup>+etoposide 200 mg/m<sup>2</sup> twice daily for 4 days+cytarabine 200 mg/m<sup>2</sup> twice daily for 4 days + single dose of melphalan 140 mg/m<sup>2</sup>; TBI+CY- 12 Gray total body irradiation in 6 fractions/cyclophosphamide 60 mg/kg daily for 2 doses (n=1); MEL- melphalan 200 mg/m<sup>2</sup>. GVHD prophylaxis: allogeneic HCT recipients received ATG 5 mg/kg in three divided doses+calcineurin inhibitors (tacrolimus or cyclosporine)+mycophenolate mofetil (15 mg/kg twice daily for 30 days) or methotrexate (5 mg/m<sup>2</sup> for four doses).

**Figure 1.**
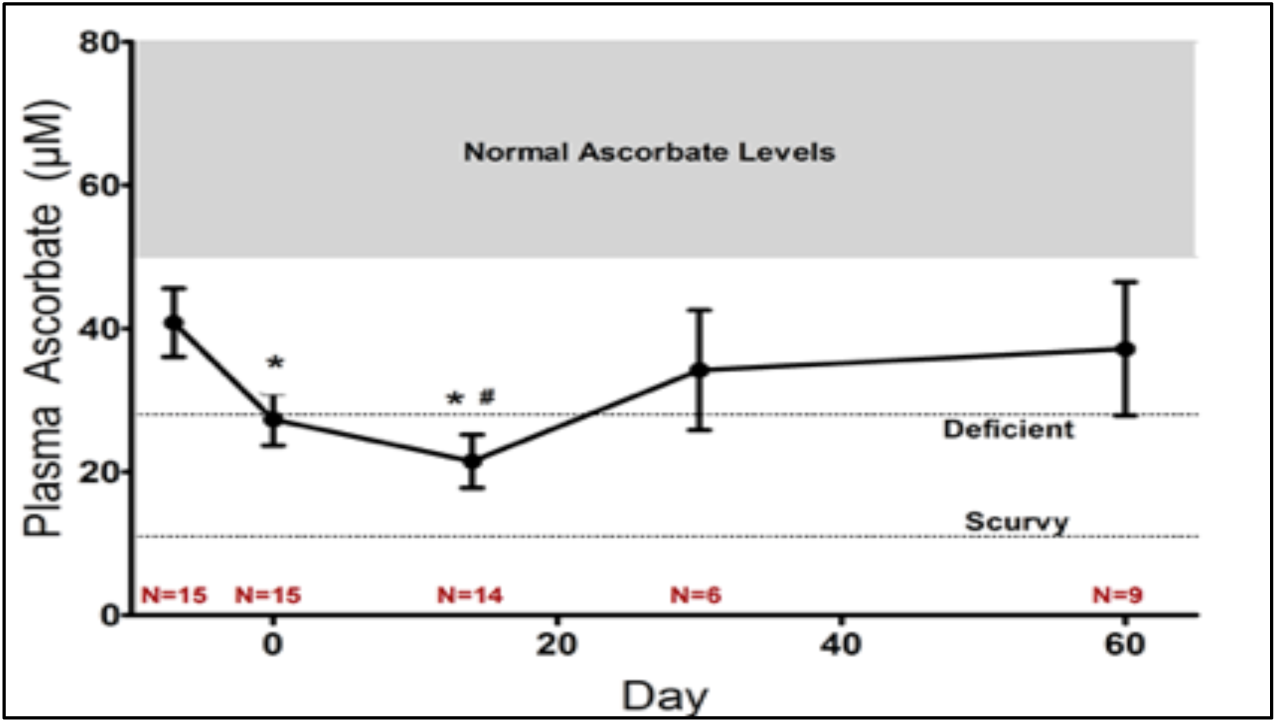
Mean plasma ascorbate levels in all transplant recipients (* P values compared to baseline). At nadir the average vitamin C levels are in the scurvy range.

No patient maintained normal ascorbic acid levels throughout the study period. Four of the fifteen patients (26.6%) recovered normal vitamin C levels by day 60. In this limited sample of patients, no association was found between type of transplant (autologous versus allogeneic), conditioning regimen, or baseline plasma vitamin C level and likelihood of recovery of normal AA levels by day 60. Patients undergoing autologous HCT maintained significantly higher mean AA levels throughout the study period compared with those undergoing allogeneic HCT. This difference was significant at baseline (mean ascorbic acid 55.9 µmol/L in the autologous SCT recipients vs. 44 in MRD, and 32.1 in the MUD SCT recipients, p < 0.05) and on day 0 (ascorbic acid 40.1 vs. 12.3 and 22.5 respectively; p < 0.05). Autologous SCT recipients also had faster post-transplant normalization of AA levels (Figure 2).

**Figure 2.**
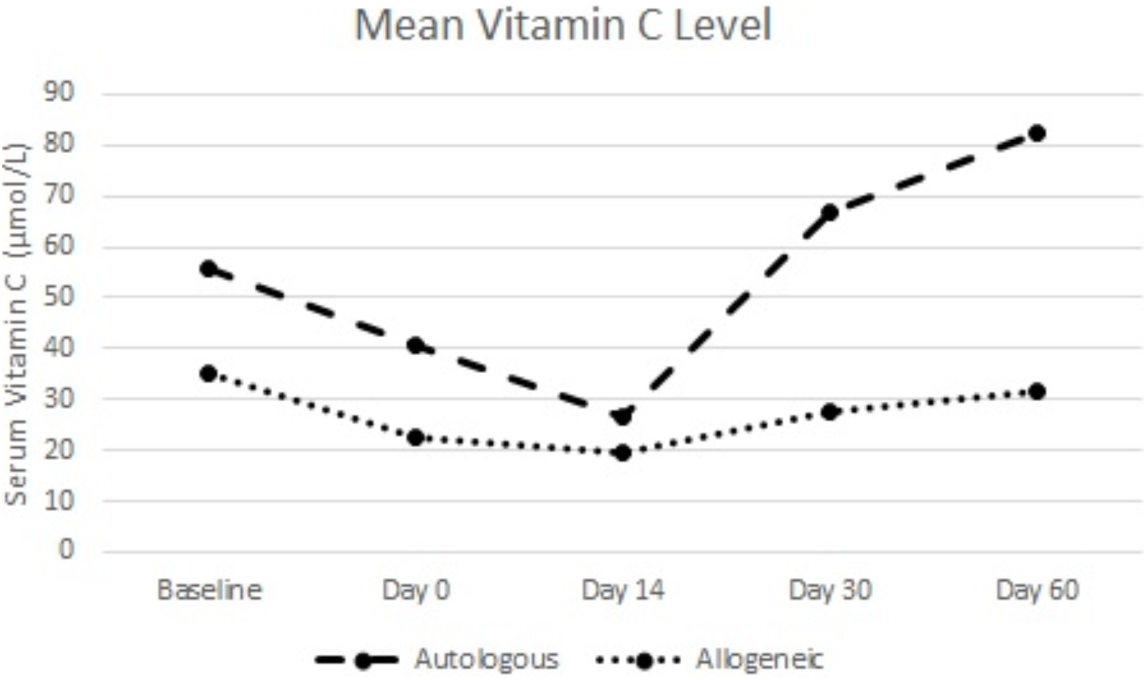
Mean plasma Ascorbic acid (Vitamin C) levels in autologous and allogeneic SCT recipients.

### Mucositis

Mucositis was recorded in 12 patients, WHO grade 1 in 3 patients, grade 2 in 5, and grade 3 in 4 patients. Severity of mucositis (determined by WHO Grade) was inversely correlated with serum ascorbic acid levels at day 14 (R −0.57, p < 0.05) (Figure 3). Of the 12 patients with mucositis post-transplant, 5 (42%) received parenteral nutrition in the post-transplant period.

**Figure 3.**
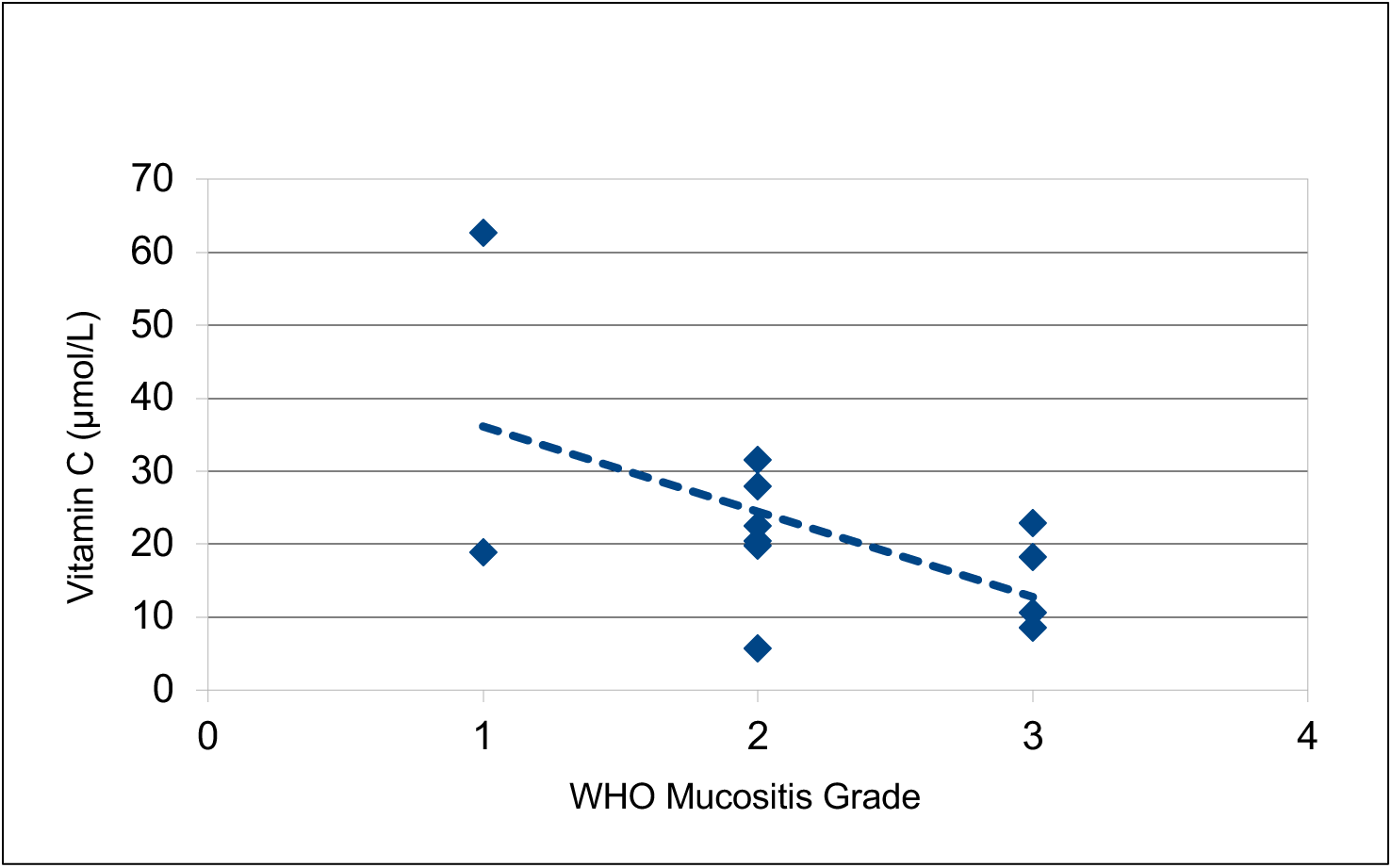
Correlation between WHO grade of mucositis and vitamin C levels on day 14

### A case of hypovitaminosis C with subsequent early lymphocyte reconstitution and GVHD

An illustrative case of development of hypovitaminosis C with subsequent early lymphocyte reconstitution and development of GVHD was observed in this cohort as follows. A 57 year-old gentleman with history of myelodysplastic syndrome, RAEB-I with complex cytogenetics, underwent allogeneic transplant from a 10/10 HLA matched unrelated donor. Busulfan, fludarabine, and antithymocyte globulin were utilized for conditioning, and tacrolimus with mycophenolate was given as GVHD prophylaxis. Baseline plasma ascorbic acid level prior to transplant was low normal at 55.2 umol/L and no mucositis was present at baseline. Grade 3 mucositis developed shortly following transplant, necessitating a continuous morphine infusion by PCA pump. Total parenteral nutrition was initiated day 6 post-transplant. Ascorbic acid plasma level at day 14 was 18.2umol/L. The patient was noted to have early lymphocyte reconstitution, with absolute lymphocyte count rising from 0 to 1200 by Day 14 with persistence of high counts (3800/µl) for the duration of the first 100 days post-transplant. Acute grade II GVHD developed ultimately required chronic, steroid, tacrolimus and extracorporeal photophoresis therapy for treatment of cutaneous, ocular, and pulmonary GHVD.

## Discussion

In this brief report a small cohort of patients undergoing myeloablative SCT who developed hypovitaminosis C following transplant is presented. No patient in this report maintained normal plasma AA levels throughout the study period. Those patients who underwent allogeneic transplant experienced lower plasma vitamin C nadirs than those undergoing autologous SCT. The severity of mucositis correlated inversely with the nadir of plasma ascorbic acid levels. The lowest ascorbic acid levels we observed was at day 14-post transplant in the immediate aftermath of most intense mucositis. This finding is similar to the observation reported earlier by Nannya and colleagues in a cohort of 15 patients, where low ascorbic acid plasma levels were observed with a nadir at day 14.^10^ Our findings extend those of Nannya et al. demonstrating that allograft recipients exhibit significant hyposcorbia several weeks into SCT despite recovery of oral intake.

Endothelial injury is a hallmark of myeloablative conditioning for SCT,^11^ with several factors such as radiation, alkylating agent exposure, and immunosuppressive drug therapy all contributing to endothelial dysfunction. In addition to prothrombotic effects, endothelial injury induces leukocyte adhesion to endothelium and increases vascular permeability. Multiple clinical manifestations ranging from pulmonary alveolar hemorrhage, capillary leak syndrome, to sinusoidal obstruction syndrome, thrombotic microangiopathy, and GVHD all may be observed. Markers of endothelial injury such as the numbers of circulating endothelial cells (CEC) increase in a dose dependent fashion following conditioning.^12^ Indeed thrombotic microangiopathy (TMA) is overrepresented in patients who develop refractory acute GVHD.^13^ Sera from GVHD patients induces expression of adhesion molecules VCAM-1, ICAM-1, and von Willebrand factor on in vitro endothelial cell cultures, suggesting an important link between endothelial dysfunction and alloreactive tissue injury.^14^

Ascorbic acid applied in the clinical appropriate clinical settings attenuates endothelial dysfunction induced by provocations such as ischemia-reperfusion injury,^15,16^ cell free hemoglobin,^17^ and sepsis.^18,19^ Ascorbic acid prevents reactive oxygen species generation through the NADPH oxidase, inducible NO synthase, and by reductive recycling of tetrahydrobiopterin.^20^ These effects occur when high doses of ascorbic acid are administered intravenously as in the sepsis trial recently reported by Fowler et al.^21^ Furthermore, high dose intravenous AA significantly reduces both vasopressor requirements as well as 28-day mortality in surgical critically ill patients.^22^ In patients with sepsis treated with intravenous AA significant reductions in C reactive protein and thrombomodulin occurred which suggests reduced inflammation and reduced vascular injury.^21^ Ascorbate is also a cofactor in the synthesis of catecholamines and vasopressin, and thus appears to have multiple critical roles in vascular biology.^23^ In animal models, parenteral AA reduces LPS induced acute lung injury and microvascular thrombosis, as well as neutrophil sequestration when administered intraperitoneally to a murine model of lung injury. Neutrophil sequestration, and lung barrier function were maintained in test animals receiving AA.^24^ In a murine model, sepsis-induced multiorgan dysfunction was attenuated by parenteral AA infusion with the mice surviving peritoneal sepsis.^25,26^

In addition to the vascular effects outlined above, it is well established that AA levels in the scurvy range (i.e., below 10 micromolar), as in our cohort may be associated with wound healing impairment.^27^ Repletion of ascorbate in these settings, accelerates wound healing and may result in attenuation of oral and gastrointestinal mucosal injury and subsequently diminished release of inflammatory cytokines (IL-1, and TNF-α), with up-regulation of mediators of wound healing (TGF-β and vascular endothelial growth factor) in animal models.^28^ Logically this will have a salutary impact on the risk of GVHD following allogeneic SCT that is classically linked to conditioning-induced tissue injury and subsequent “cytokine storm”.^29^ This raises the question of whether patients post HCT may benefit from repletion of AA, which may in turn may diminish the severity of mucositis and possibly ameliorate GVHD risk. A prospective clinical trial of parenteral ascorbic acid repletion in allograft recipients is underway (NCT03613727) to investigate this question. The logic for use in this setting is supported by preliminary data from a parenteral AA repletion study, where AA was administered to patients with severe sepsis and low serum ascorbic acid levels.^18^ In this study, the effects of intravenous ascorbic acid on inflammation and organ failure in severe sepsis were studied using the sequential organ failure scores (SOFA), and the inflammatory biomarker CRP, which declined rapidly in patients who received intravenous ascorbic acid compared to placebo. The pathophysiology induced by sepsis, with increased pro-inflammatory cytokines and endothelial damage is analogous to the *regimen-related toxicity* induced by myeloablative conditioning in HCT. These data as presented here suggest that intravenous ascorbic acid repletion may similarly benefit allograft recipients who are ascorbic acid deficient. Inflammatory conditions in early phases post HCT profoundly influence the risk of GVHD.^30,31^ Thus a non-immunosuppressive, anti-inflammatory agent such as AA may ameliorate the risk of GVHD and enhance normal T cell differentiation.

There is evidence that AA has a role in promoting the development of T cells. This phenomenon may be of critical importance in immune reconstitution following HCT. ^10^ While AA has a significant role in immune function,^32^ it has several well-characterized effects on T cells, (e.g., diminish apoptosis^33^) as well as its crucial impact on the development of double positive and single positive, T cell receptor-rearranged, T cells in feeder-free cell culture systems.^34^ In particular, the effects on CD8 and ZAP-70 as well as IL-17 expression appear to be mediated via chromatin methylation through the Jomanji C domain enzyme that employs ascorbate as a cofactor for promoting histone demethylation.^35,36^ Vitamin C has thus been suggested as a mediator of enhanced immune reconstitution.^37^ Indeed, in a GVHD murine model, vitamin C stabilizes alloreactive T regulatory cells by demethylation of the FoxP3 promoter, the intronic cis-regulatory elements on DNA molecules.^38^ Ascorbic acid is also a cofactor for the enzyme ten-eleven translocation (TET) which mediates promoter hypomethylation of the FOX P3 locus amongst other loci involved in hematopoiesis.^39,40^ Huijskens et al. showed that AA enhances ex vivo expansion of NK cells in cell culture systems as well as their differentiation from hematopoietic progenitors.^41^ A critically important recent discovery is that AA controls leukemic cell proliferation and differentiation, particularly in TET2 and IDH1 mutated AML. This effect is mediated through epigenetic control of transcription factor binding sites in leukemia.^42,43,44,45^ These results suggest that post-transplant AA repletion may contribute to the control of hematological malignancies.

In conclusion, AA exhibits pleiotropic effects in human physiology. These effects may broadly impact clinical outcomes in SCT recipients. Myeloablative allogeneic stem cell transplant recipients are deficient in ascorbic acid. Patients receiving SCT may benefit from intravenous AA repletion, particularly in the early weeks following transplant. Studies aimed at repletion of AA in allogeneic HCT recipients to determine the effects on endothelial dysfunction and immune reconstitution are critical for development of strategies to thwart the dual threat of transplant related mortality and relapse.

## Acknowledgement

Funding for this study was provided by Massey Cancer Center Pilot Project Grant.

## Authorship Contributions

Mahmood Rasheed collected and analyzed data and wrote the paper. Gary L. Simmons analyzed data and wrote the paper. Bernard J. Fisher contributed vital analytical tools and performed analysis of plasma AA levels. Cathy Roberts analyzed clinical data. Alpha A. Fowler designed the research. Ramesh Natarajan contributed vital analytical tools, performed research and wrote the paper. Amir Toor designed research and wrote the paper. AAT was supported, by research funding from the NIH-NCI Cancer Center Support Grant (P30-CA016059; PI: Gordon Ginder, MD).

